# Dynamics of scene representations in the human brain revealed by magnetoencephalography and deep neural networks

**DOI:** 10.1101/032623

**Authors:** Radoslaw Martin Cichy, Aditya Khosla, Dimitrios Pantazis, Aude Oliva

**Affiliations:** Computer Science and Artificial Intelligence Laboratory, MIT, Cambridge, MA, USA; McGovern Institute for Brain Research, MIT, Cambridge, MA, USA

**Keywords:** Scene perception, spatial layout, magnetoencephalography, deep neural network, representational similarity analysis

## Abstract

Human scene recognition is a rapid multistep process evolving over time from single scene image to spatial layout processing. We used multivariate pattern analyses on magnetoencephalography (MEG) data to unravel the time course of this cortical process. Following an early signal for lower-level visual analysis of single scenes at ~100ms, we found a marker of real-world scene size, i.e. spatial layout processing, at ~250ms indexing neural representations robust to changes in unrelated scene properties and viewing conditions. For a quantitative explanation that captures the complexity of scene recognition, we compared MEG data to a deep neural network model trained on scene classification. Representations of scene size emerged intrinsically in the model, and resolved emerging neural scene size representation. Together our data provide a first description of an electrophysiological signal for layout processing in humans, and a novel quantitative model of how spatial layout representations may emerge in the human brain.

## 1 INTRODUCTION

Perceiving the geometry of space is a core ability shared by all animals, with brain structures for spatial layout perception and navigation preserved across rodents, monkeys and humans (Epstein and Kanwisher, 1998, 1998; Doeller et al., 2008, 2010; Moser et al., 2008; Epstein, 2011; Jacobs et al., 2013; Kornblith et al., 2013, 2013; Vaziri et al., 2014). Spatial layout perception, the demarcation of the boundaries and size of real-world visual space, plays a crucial mediating role in spatial cognition (Bird et al., 2010; Epstein, 2011; Kravitz et al., 2011a; Wolbers et al., 2011a; Park et al., 2014) between image-specific processing of individual scenes and navigation-related processing. Although the cortical loci of spatial layout perception in humans have been well described (Aguirre et al., 1998; Kravitz et al., 2011b; MacEvoy and Epstein, 2011; Mullally and Maguire, 2011; Park et al., 2011; Bonnici et al., 2012), the dynamics of spatial cognition remain unexplained, partly because neuronal markers indexing spatial processing remain unknown.

Operationalizing spatial layout as scene size, that is the size of the space a scene subtends in the real-world (Kravitz et al., 2011a; Park et al., 2011, 2014), we report here an electrophysiological signal of spatial layout perception in the human brain. Using multivariate pattern classification (Carlson et al., 2013; Cichy et al., 2014; Isik et al., 2014) and representational similarity analysis (Kriegeskorte, 2008; Kriegeskorte and Kievit, 2013; Cichy et al., 2014) on millisecond-resolved magnetoencephalography data (MEG), we identified a marker of scene size around 250ms, preceded by and distinct from an early signal for lower-level visual analysis of scene images at ~100ms.

Furthermore, we demonstrated that the scene size marker was independent of both low-level image features (i.e. luminance, contrast, clutter) and semantic properties (the category of the scene, i.e. kitchen, ballroom), thus indexing neural representations robust to changes in viewing conditions as encountered in real-world settings.

To provide a quantitative explanation how space size representations emerge in cortical circuits, we compared brain data to a deep neural network model trained to perform scene categorization (Zhou et al., 2014, 2015), termed deep scene network. The deep scene network *intrinsically* exhibited receptive fields specialized for layout analysis, such as textures and surface layout information, without ever having been explicitly taught any of those features. We showed that the deep scene neural network model predicted the human neural representation of single scenes and scene space size better than a deep object model and standard models of scene and object perception (Riesenhuber and Poggio, 1999; Oliva and Torralba, 2001). This demonstrates the ability of the deep scene model to approximate human neural representations at successive levels of processing as they emerge over time.

Together our findings provide a first description of an electrophysiological signal for scene space processing in humans, and offer a novel quantitative and computational model of the dynamics of visual scene space representation in the cortex. Our results suggest that spatial layout representations naturally emerge in cortical circuits learning to differentiate visual environments (Oliva and Torralba, 2001).

## 2 MATERIALS AND METHODS

### 2.1 Participants

Participants were 15 right-handed, healthy volunteers with normal or corrected-to-normal vision (mean age ± s.d. = 25.87 ± 5.38 years, 11 female). The Committee on the Use of Humans as Experimental Subjects (COUHES) at MIT approved the experiment and each participant gave written informed consent for participation in the study, for data analysis and publication of study results.

### 2.2 Stimulus material and experimental design

The image set consisted of 48 scene images differing in four factors with two levels each, namely two scene properties: physical size (small, large) and clutter level (low, high); and two image properties: contrast (low, high) and luminance (low, high) (Figure 1A). There were 3 unique images for every level combination, for example 3 images of small size, low clutter, low contrast and low luminance. The image set was based on behaviorally validated images of scenes differing in size and clutter level, sub-sampling the two highest and lowest levels of factors size and clutter (Park et al., 2014). Small scenes were of size that would typically fit 2-8 people, whereas large scenes would fit hundreds to thousands. Similarly, low clutter level scenes were empty or nearly empty rooms, whereas high clutter scenes contained multiple objects throughout. The contrast and luminance was adjusted to specific values for each image: images of low and high contrast had root mean square values of 34% and 50% respectively; images of low and high luminance had root mean square values of 34% and 51% respectively.

**Figure 1.**
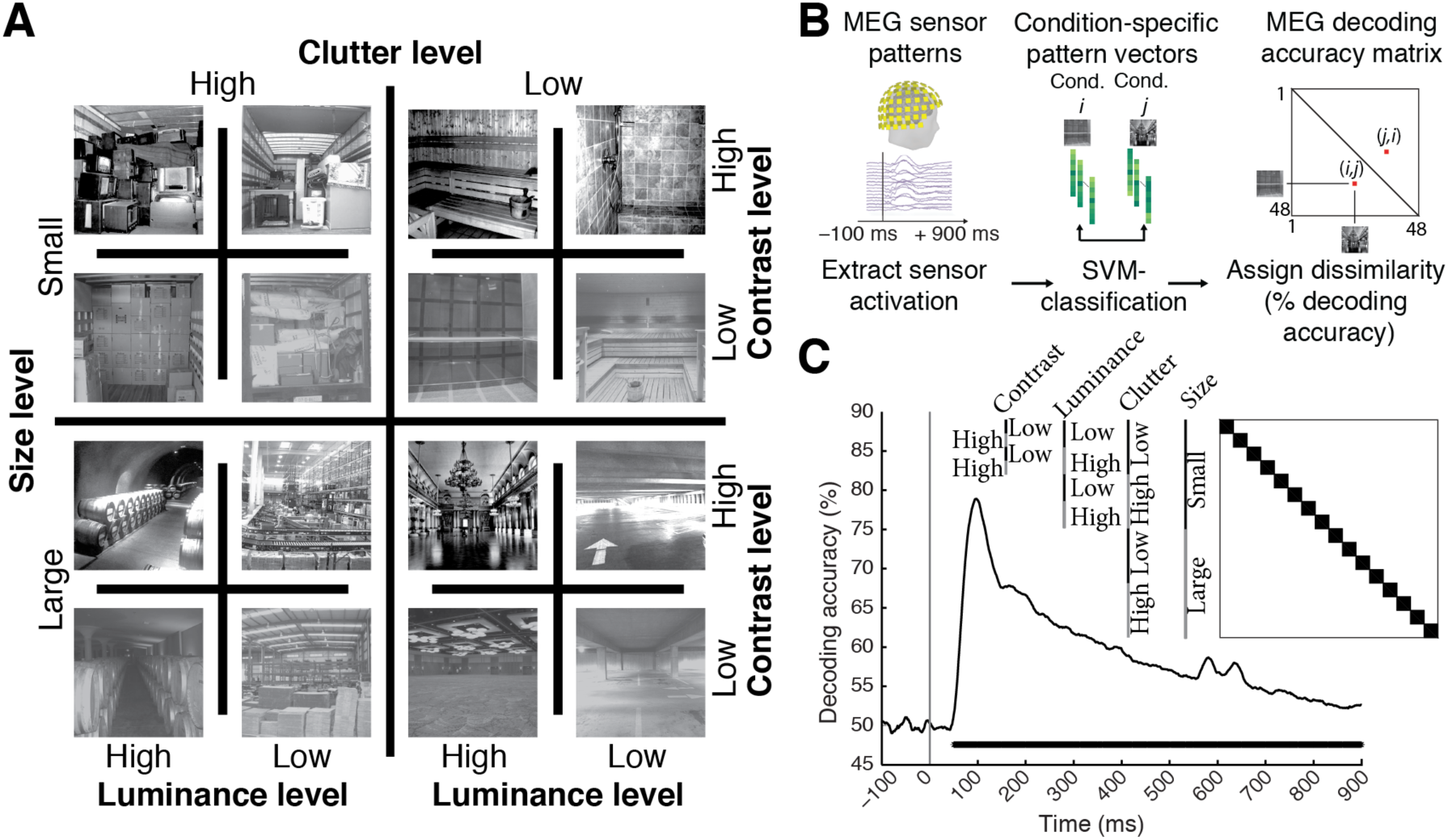
Image set and single-image decoding. **A**) The stimulus set comprised 48 indoor scene images differing in the size of the space depicted (small vs. large), as well as clutter, contrast, and luminance level; here each experimental factor combination is exemplified by one image. The image set was based on behaviorally validated images of scenes differing in size and clutter level, de-correlating factors size and clutter explicitly by experimental design (Park et al., 2014). Note that size refers to the size of the real-world space depicted on the image, not the stimulus parameters; all images subtended 8 visual angle during the experiment. **B**) Time-resolved (1ms steps from -100 to +900ms with respect to stimulus onset) pair-wise support vector machine classification of experimental conditions based on MEG sensor level patterns. Classification results were stored in time-resolved 48 × 48 MEG decoding matrices. **C**) Decoding results for single scene classification independent of other experimental factors. Decoding results were averaged across the dark blocks (matrix inset), to control for luminance, contrast, clutter level and scene size differences. Inset shows indexing of matrix by image conditions. Horizontal line below curve indicates significant time points (*n* = 15, cluster-definition threshold *P* < 0.05, corrected significance level *P* < 0.05); gray vertical line indicates image onset.

Participants viewed a series of scene images while MEG data was recorded (Figure 1B). Images subtended 8° of visual angle in both width and height and were presented centrally on a gray screen (42.5% luminance) for 0.5s in random order with an inter-stimulus interval (ISI) of 1-1.2s, overlaid with a central red fixation cross. Every 4 trials on average (range 3-5 trials, equally probable) a target image depicting concentric circles was presented prompting participants to press a button and blink their eyes in response. ISI between the concentric-circles and the next trial was 2s to allow time for eye blinks. Target image trials were not included in analysis. Each participant completed 15 runs of 312s each. Every image was presented four times in a run, resulting in 60 trials per image per participant in total.

### 2.3 MEG recording

We recorded continuous MEG signals from 306 channels (Elektra Neuromag TRIUX, Elekta, Stockholm) at a sampling rate of 1000Hz. Raw data was band-pass filtered between 0.03 and 330Hz, and pre-processed using spatiotemporal filters (maxfilter software, Elekta, Stockholm). We used Brainstorm (Tadel et al., 2011) to extract peristimulus MEG signals from −100 to +900ms with respect to stimulus onset, and then normalized each channel by its baseline (−100 to 0ms) mean and standard deviation, and temporally smoothed the time series with a 20ms sliding window.

### 2.4 Multivariate pattern classification of MEG data

*Single image classification:* To determine whether MEG signals can discriminate experimental conditions (scene images), data were subjected to classification analyses using linear support-vector machines (SVM) (Müller et al., 2001) in the libsvm implementation (www.csie.ntu.edu.tw/~cjlin/libsvm) with a fixed regularization parameter C=1. For each time point t, the processed MEG sensor measurements were concatenated to 306-dimensional pattern vectors, resulting in M=60 raw pattern vectors per condition (Figure 1B). To reduce computational load and improve signal-to-noise ratio, we sub-averaged the M vectors in groups of k = 5 with random assignment, thus obtaining M/k averaged pattern vectors. We then measured the performance of the SVM classifier to discriminate between every pair (i,j) of conditions using a leave-one-out approach: M/k - 1 vectors were randomly assigned to the training test, and 1 vector to the testing set to evaluate the classifier decoding accuracy. The above procedure was repeated 100 times, each with random assignment of the M raw pattern vectors to M/k averaged pattern vectors, and the average decoding accuracy was assigned to the (i,j) element of a 48 x 48 decoding matrix indexed by condition. The decoding matrix is symmetric with an undefined diagonal. We obtained one decoding matrix (representational dissimilarity matrix or RDM) for each time point t.

*Representational clustering analysis for size:* Interpreting decoding accuracy as a measure of dissimilarity between patterns, and thus as a distance measure in representational space (Kriegeskorte and Kievit, 2013; Cichy et al., 2014), we partitioned the RDM decoding matrix into within- and between-level segments for the factor scene size (Figure 2A). The average of between-size minus within-size matrix elements produced representational distances (percent decoding accuracy difference) indicative of clustering of visual representations by scene size.

**Figure 2.**
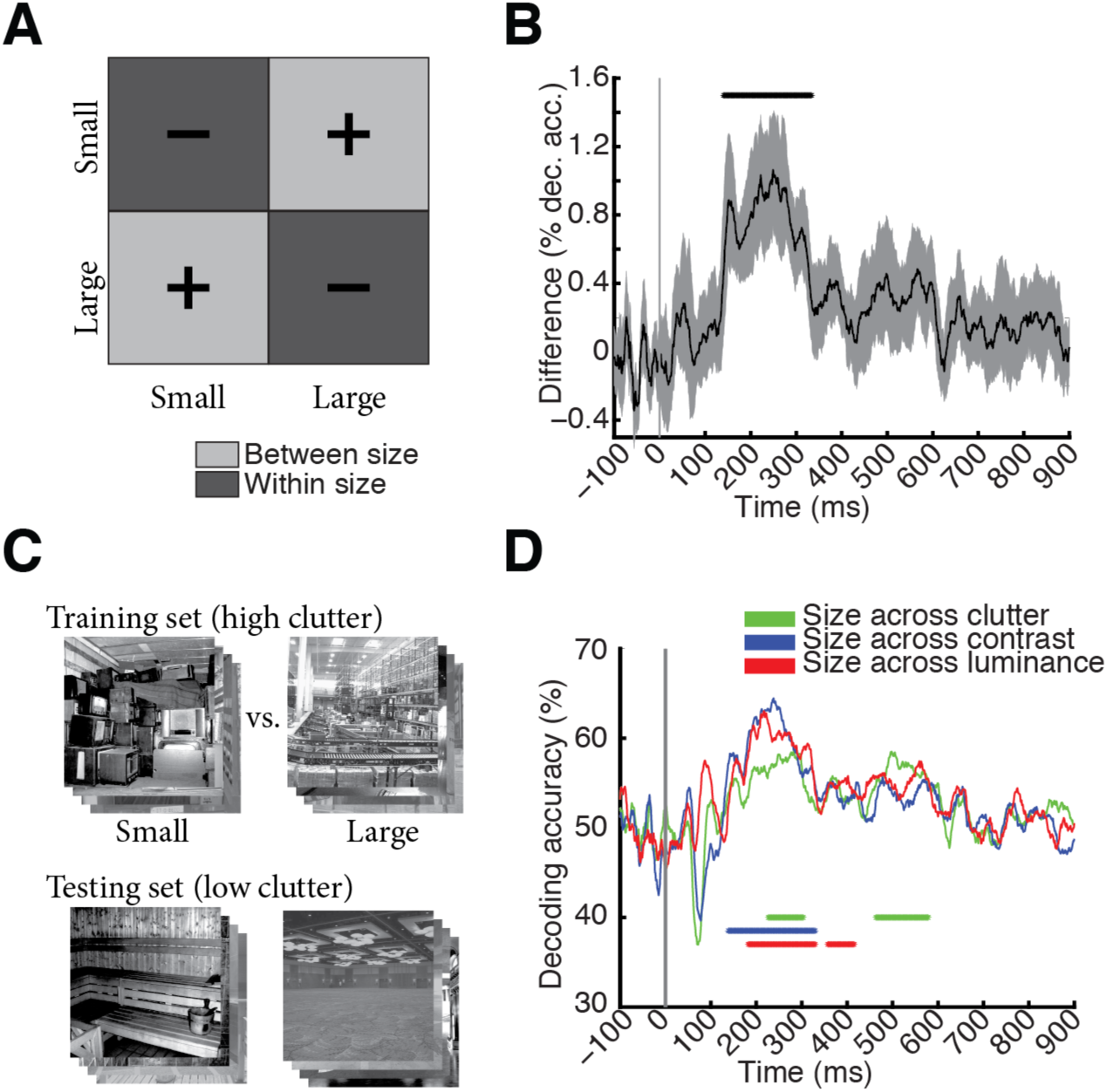
Scene size is discriminated by visual representations. **A)** To determine the time course of scene size processing we determined when visual representations clustered by scene size. For this we subtracted mean within-size decoding accuracies (dark gray, −) from between-size decoding accuracies (light gray, +). **B)** Scene size was discriminated by visual representations late in time (onset of significance at 141ms (118-156ms), peak at 249ms (150-274ms). Gray shaded area indicates 95% confidence intervals determined by bootstrapping participants. **C)** Cross-classification analysis, exemplified for cross-classification of scene size across clutter level. A classifier was trained to discriminate scene size on high clutter images, and tested on low clutter images. Results were averaged following an opposite assignment of clutter images to training and testing sets. Before entering cross-classification analysis, MEG trials were grouped by clutter and size level respectively independent of image identity. A similar cross-classification analysis was applied for other image and scene properties. **D)** Results of cross-classification analysis indicated robustness of scene size visual representations to changes in other scene and image properties (scene clutter, luminance, and contrast). Horizontal lines indicate significant time points *(n* = 15, cluster-definition threshold *P* < 0.05, corrected significance level *P* < 0.05); gray vertical line indicates image onset. For result curves with 95% confidence intervals see Supplementary Figure 2.

*Cross-classification:* To assess whether scene size representations were robust to changes of other factors, we used SVM cross-classification assigning different levels of experimental factors to the training and testing set. For example, Figure 2C shows the cross-classification of scene size (small vs. large) across clutter, implemented by limiting the training set to high clutter scenes and the testing set to low clutter scenes. The procedure was repeated with reverse assignment (low clutter for training set and high clutter for testing set) and decoding results were averaged. The training set was 12 times larger (*M* = 720 raw pattern vectors) than for single-image decoding, as we pooled trials across single images that had the same level of clutter and size. We averaged pattern vectors by sub-averaging groups of *k* = 60 raw pattern vectors before the leave-one-out SVM classification. Cross-classification analysis was performed for the cross-classification of the factors scene size (Figure 2D) and scene clutter (Supplementary Figure 3) with respect to changes across all other factors.

### 2.5 Low and high-level computational models of image statistics

We assessed whether computational models of object and scene recognition predicted scene size from our image material. For this we compared four models: two deep convolutional neural networks that were either trained to perform (1) scene or (2) object classification; (3) the GIST descriptor (Oliva and Torralba, 2001), i.e. a model summarizing the distribution of orientation and spatial frequency in an image that has been shown to predict scene properties, among them size; and (4) HMAX model (Serre et al., 2005), a model of object recognition most akin in structure to low-level visual areas V1/V2. We computed the output of each of these models for each image as described below.

#### Deep neural networks

The deep neural network architecture was implemented following Krizhevsky et al., 2012. We chose this particular architecture because it was the best performing model in object classification in the ImageNet 2012 competition (Russakovsky et al., 2014), uses biologically-inspired local operations (convolution, normalization, max-pooling), and has been compared to human and monkey brain activity successfully (Güçlü and van Gerven, 2014; Khaligh-Razavi and Kriegeskorte, 2014; Khaligh-Razavi et al., 2014). The network architecture had 8 layers with the first 5 layers being convolutional and the last 3 fully connected. For an enumeration of units and features for each layer see Table 3. We used the convolution stage of each layer as model output for further analysis.

We constructed two deep neural networks that differed in the visual categorization task and visual material they were trained on. A deep scene model was trained on 216 scene categories from the Places dataset (available online at: http://places.csail.mit.edu/) (Zhou et al., 2015) with 1300 images per category. A deep object model was trained on 683 different objects with 900,000 images from the ImageNet dataset (available online at: http://www.image-net.org/) (Deng et al., 2009) with similar number of images per object category (~1300). Both deep neural networks were trained on GPUs using the Caffe toolbox (Jia et al., 2014). In detail, the networks were trained for 450,000 iterations, with an initial learning rate of 0.01 and a step multiple of 0.1 every 100,000 iterations. Momentum and weight decay were kept constant at 0.9 and 0.0005 respectively.

To visualize receptive fields (RFs) of model neurons in the deep scene network (Figure 3B) we used a reduction method (Zhou et al., 2015). In short, for a particular neuron we determined the *K* images activating the neuron most strongly. To determine the empirical size of the RF, we replicated the K images many times with small random occluders at different positions in the image. We then passed the occluded images into the deep scene network and compared the output to the original image, constructing the discrepancy map that indicates which part of the image drives the neuron. We then recentered discrepancy maps and averaged, generating the final RF. To illustrate the RFs tuning we further plot the image patches corresponding to the top activation regions inside the RFs (Figure 3B).

**Figure 3.**
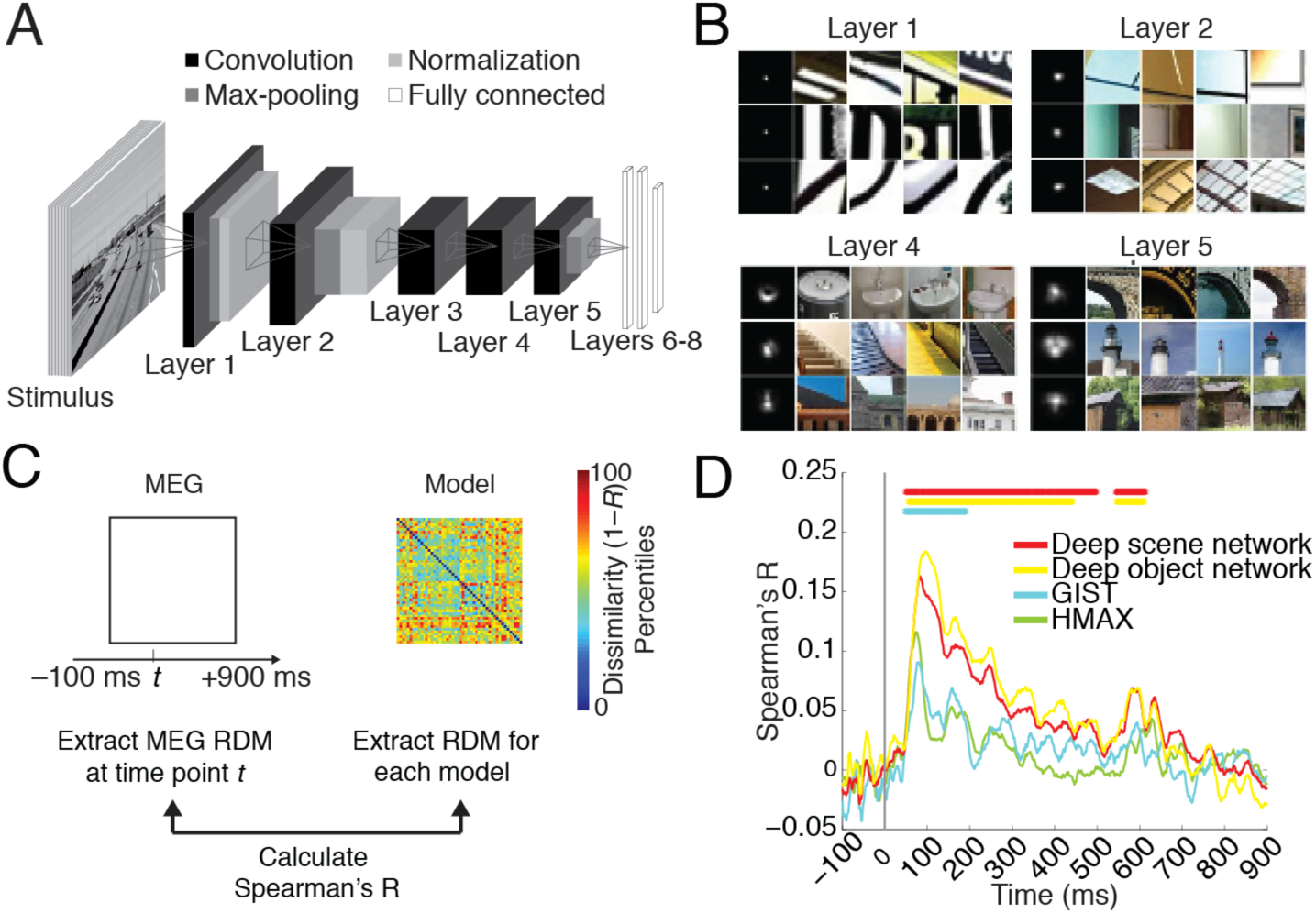
Predicting emerging neural representations of single scene images by computational models. **A)** Architecture of deep convolutional neural network trained on scene categorization (deep scene network). **B)** Receptive field (RF) of example deep scene neurons in layers 1, 2, 4, and 5. Each row represents one neuron. The left column indicates size of RF, and the remaining columns indicate image patches most strongly activating these neurons. Lower layers had small RFs with simple Gabor filter-like sensitivity, whereas higher layers had increasingly large RFs sensitive to complex forms. RFs for whole objects, texture, and surface layout information emerged although these features were not explicitly taught to the deep scene model. **C)** We used representational dissimilarity analysis to compare visual representations in brains with models. For every time point, we compared subject-specific MEG RDMs (Spearman’s R) to model RDMs and results were averaged across subjects. **D)** All investigated models significantly predicted emerging visual representations in the brain, with superior performance for the deep neural networks compared to HMAX and GIST. Horizontal lines indicate significant time points *(n* = 15, cluster-definition threshold *P* < 0.05, corrected significance level *P* < 0.05); gray vertical line indicates image onset.

#### GIST

For the GIST descriptor (Oliva and Torralba, 2001), each image was filtered by a bank of Gabor filters with 8 orientations and 4 spatial frequencies (32 filters). Filter outputs were averaged in a 4×4 grid, resulting in a 512-dimensional feature vector. The GIST descriptor represents images in terms of spatial frequencies and orientations by position, (code available: *http://people.csail.mit.edu:/torralba/code/spatialenvelope/)*.

#### HMAX

We used the HMAX model as applied and described by Serre et al (Serre et al., 2005), a model inspired by the hierarchical organization of the visual cortex. In short, HMAX consists of two sets of alternating S and C layers, i.e. in total 4 layers. The S-layers convolve the input with pre-defined filters, and the C layers perform a max operation.

### 2.6 Linking computational models of vision to brain data

We used representational dissimilarity analysis to compare the output of computational models to brain data. First, we recorded the output of each model for each of the 48 images of the image set. Then, to compare to human brain data, we calculated the pair-wise dissimilarities between model outputs by 1-Spearman’s rank order correlation *R*. This formed 48x48 model dissimilarity matrices (RDMs), one for each layer of each model: 8 for the deep scene and deep object network, 1 for GIST, and 4 for HMAX.

To compare models and brains, we determined whether images that were similarly represented in a computational network were also similarly represented in the brain. This was achieved by computing the similarity (Spearman’s *R*) of layer-specific model dissimilarity matrix with the time-point specific MEG decoding matrix for every subject and time point and averaging results.

We then determined whether the computational models predicted the size of a scene. We formulated an explicit size model, i.e. a 48 × 48 matrix with entries of 1 where images differed in size and 0 otherwise. Equivalent matrices were produced for scene clutter, contrast and luminance (Supplementary Figure 1). Correlation of the explicit size model with any computational model RDM yielded a measure of how well computational models predicted scene size.

Finally, we determined whether the above computational models accounted for neural representations of scene size observed in MEG data. For this, we reformulated the representational clustering analysis in a correlation framework. The two measures are equivalent except that the correlation analysis takes into account the variability of the data, which the clustering analysis does not for the benefit of clear interpretability as percent change in decoding accuracy. The procedure had two steps. First, we calculated the similarity (Spearman’s *R)* of the MEG decoding accuracy matrix with the explicit size model for each time point and each participant. Second, we re-calculated the similarity (Spearman’s *R)* of the MEG decoding accuracy matrix with the explicit size model after partialling out all of the layer-specific RDMs of a given computational model.

### 2.7 Statistical testing

We used permutation tests for cluster-size inference, and bootstrap tests to determine confidence intervals of onset times for maxima, cluster onsets and peak-to-peak latency differences (Nichols and Holmes, 2002; Pantazis et al., 2005; Cichy et al., 2014).

#### Sign permutation tests

For the permutation tests, depending on the statistic of interest our null hypothesis was that the MEG decoding time series were equal to 50% chance level, or that the decoding accuracy difference of between-minus within-level segments of the MEG decoding matrix was equal to 0, or that the correlation values were equal to 0. In all cases, under the null hypothesis the sign of the observed effect in the MEG data is randomly permutable, corresponding to a sign-permutation test that randomly multiplies the participant-specific data with +1 or −1. We created 1,000 permutation samples, every time re-computing the statistic of interest. This resulted in an empirical distribution of the data, allowing us to convert our original data, as well as the permutation samples, into *P*-values. We then performed cluster-size inference by setting a *P* = 0.05 cluster-definition threshold on the original data and permutation samples, and computing a *P* = 0.05 cluster size threshold from the empirical distribution of the resampled data.

#### Bootstrapping

To calculate confidence intervals (95%) on cluster onset and peak latencies, we bootstrapped the sample of participants 1,000 times with replacement. For each bootstrap sample, we repeated the above permutation analysis yielding distributions of the cluster onset and peak latency, allowing estimation of confidence intervals. In addition, for each bootstrap sample, we determined the peak-to-peak latency difference for scene size clustering and individual scene image classification. This yielded an empirical distribution of peak-to-peak latencies. Setting *P* < 0.05, we rejected the null hypothesis of a latency difference if the confidence interval did not include 0.

#### Label permutation tests

For testing the significance of correlation between the computational model RDMs and the scene size model, we relied on a permutation test of image labels. This effectively corresponded to randomly permuting the columns (and accordingly the rows) of the computational model RDMs 1,000 times, and then calculating the correlation between the permuted matrix and the explicit size model matrix. This yielded an empirical distribution of the data, allowing us to convert our statistic into *P*-values. Effects were reported as significant when passing a *P* = 0.05 threshold. Results were FDR-corrected for multiple comparisons.

## 3 RESULTS

Human participants (*n* = 15) viewed images of 48 real-world indoor scenes that differed in the layout property size, as well as in the level of clutter, contrast and luminance (Figure 1A) while brain activity was recorded with MEG. While often real-world scene size and clutter level correlate, here we de-correlated those stimulus properties explicitly by experimental design, based on independent behavioral validation (Park et al., 2014) to allow independent assessment. Images were presented for 0.5s with an inter-trial interval of 1-1.2s (Figure 1B). Participants performed an orthogonal object-detection task on an image of concentric circles appearing every four trials on average. Concentric circle trials were excluded from further analysis.

To determine the timing of cortical scene processing we used a decoding approach: we determined the time course with which experimental conditions (scene images) were discriminated by visual representations in MEG data. For this, we extracted peri-stimulus MEG time series in 1ms resolution from -100 to +900ms with respect to stimulus onset for each subject. For each time point independently we classified scene images pair-wise by MEG sensor patterns (support vector classification, Figure 1C). Time-point specific classification results (percentage decoding accuracy, 50% chance level) were stored in a 48×48 decoding accuracy matrix, indexed by image conditions in rows and columns (Figure 1C, inset). This matrix is symmetric with undefined diagonal. Repeating this procedure for every time point yielded a set of decoding matrices (for a movie of decoding accuracy matrices over time, averaged across subjects, see Supplementary Movie 1). Interpreting decoding accuracies as a representational dissimilarity measure, each 48x48 matrix summarized, for a given time point, which conditions were represented similarly (low decoding accuracy) or dissimilarly (high decoding accuracy). The matrix was thus termed MEG representational dissimilarity matrix (RDM) (Cichy et al., 2014; Nili et al., 2014).

Throughout, we determined random-effects significance non-parametrically using a cluster-based randomization approach (cluster-definition threshold *P* < 0.05, corrected significance level *P* < 0.05) (Nichols and Holmes, 2002; Pantazis et al., 2005; Maris and Oostenveld, 2007). 95% confidence intervals for mean peak latencies and onsets (reported in parentheses throughout the results) were determined by bootstrapping the participant sample.

### 3.1 Neural representations of single scene images emerged early in cortical processing

We first investigated the temporal dynamics of image-specific individual scene information in the brain. To determine the time course with which individual scene images were discriminated by visual representations in MEG data, we averaged the elements of each RDM matrix representing pairwise comparisons with matched experimental factors (luminance, contrast, clutter level and scene size) (Figure 1C). We found that the time course rose sharply after image onset, reaching significance at 50ms (45-52ms) and a peak at 97ms (94-102ms). This indicates that single scene images were discriminated early by visual representations, similar to single images with other visual content (Thorpe et al., 1996; Carlson et al., 2013; Cichy et al., 2014; Isik et al., 2014), suggesting a common source in early visual areas (Cichy et al., 2014).

### 3.2 Neural representations of scene size emerged later in time and were robust to changes in viewing conditions and other scene properties

When is the spatial layout property scene size processed by the brain? To investigate, we partitioned the decoding accuracy matrix into two subdivisions: images of different (between subdivision light gray, +) and similar size level (within subdivision, dark gray, −). The difference of mean between-size minus within-size decoding accuracy is a measure of clustering of visual representations by size (Figure). Peaks in this measure indicate time points at which MEG sensor patterns cluster maximally by scene size, suggesting underlying neural visual representations allowing for explicit, linear readout (DiCarlo and Cox, 2007) of scene size by the brain. Scene size (Figure 2B) was discriminated first at 141ms (118 – 156ms) and peaked at 249ms (150 – 274ms), which was significantly later than the peak in single image classification (*P* = 0.001, bootstrap test of peak-latency differences).

Equivalent analyses for the experimental factors scene clutter, contrast, and luminance level yielded diverse time courses (Supplementary Figure 1, Table 1A). Importantly, representations of low-level image property contrast emerged significantly earlier than scene size (*P* = 0.004) and clutter (*P* = 0.006, bootstrap test of peak-latency differences). For the factor luminance, only a weak effect and thus no significant onset response was observed, suggesting a pre-cortical luminance normalization mechanism.

**Table 1.** Onset and peak latencies for MEG classification analyses. Onset and peak latency *(n* = 15, *P* < 0.05, cluster-level corrected, cluster-definition threshold *P* < 0.05) with 95% confidence intervals. **A**) Clutter, luminance and contrast level representation time course information. **B**) Time course of cross-classification for scene size. 95% confidence intervals are reported in brackets.

| **A** |  |  |
| --- | --- | --- |
|  | **Onset latency** | **Peak latency** |
| Clutter level | 56 (42 – 71) | 107 (103 – 191) |
| Luminance level | 644 (68 – 709) | 625 (146 – 725) |
| Contrast level | 53 (42 – 128) | 74 (68 – 87) |
| **B** |  |  |
| Size across clutter level | 226 (134 – 491) | 283 (191 – 529) |
| Size across luminance level | 183 (138 – 244) | 217 (148 – 277) |
| Size across contrast level | 138 (129 – 179) | 238 (184 – 252) |

To be of use in the real world, visual representations of scene size must be robust against changes of other scene properties, such as clutter level (i.e. space filled by different types and amounts of objects) and semantic category (i.e. the label by which we name it), and changes in viewing conditions, such as luminance and contrast. We investigated the robustness of scene size representations to all these factors using cross-classification (Figure 2C; for 95% confidence intervals on curves see Supplementary Figure 2). For this we determined how well a classifier trained to distinguish scenes at one clutter level could distinguish scenes at the other level, while collapsing data across single image conditions of same level in size and clutter. We found that scene size was robust to changes in scene clutter, luminance and contrast (Figure 2D; onsets and peaks in Table 1B). Note that by experimental design, the scene category always differed across size level, such that cross-classification also established that scene size was discriminated by visual representations independent of the scene category.

An analogous analysis for clutter level yielded evidence for viewing-condition independent clutter level representations (Supplementary Figure 3), reinforcing the notion of clutter level as a robust and relevant dimension of scene representations in the human brain (Park et al., 2014). Finally, an analysis revealing persistent and transient components of scene representations indicated strong persistent components for scene size and clutter representations, with little or no evidence for contrast and luminance (Supplementary Figure 4). Persistence of scene size and clutter level representations further reinforces the notion of size and clutter level representations being important end products of visual computations kept online by the brain for further processing and behavioral guidance.

In sum, our results constitute evidence for representations of scene size in human brains from non-invasive electrophysiology, apt to describe scene size discrimination under real world changes in viewing conditions.

### 3.3 Neural representations of single scene images were predicted by deep convolutional neural networks trained on real world scene categorization

Visual scene recognition in cortex is a complex hierarchical multi-step process, whose understanding necessitates a quantitative model that captures this complexity. Here, we evaluated whether an 8-layer deep neural network trained to perform scene classification on 205 different scene categories (Zhou et al., 2014) predicted human scene representations. We refer to this network as deep scene network (Figure 3A). Investigation of the receptive fields (RFs) of model neurons using a reduction method (Zhou et al., 2015) indicated a gradient of increasing complexity from low to high layers, and selectivity to whole objects, texture, and surface layout information (Figure 3B). This suggests that the network might be able to capture information about both single scenes and scene layout properties.

To determine the extent to which visual representations learned by the deep scene model and the human brain are comparable, we used representational similarity analysis (Kriegeskorte, 2008; Cichy et al., 2014). The key idea is that if two images evoke similar responses in the model, they should evoke similar responses in the brain, too.

For the deep neural network, we first estimated image response patterns by computing the output of each model layer to each of the 48 images. We then constructed layer-resolved 48×48 representational dissimilarity matrices (RDMs) by calculating the pairwise dissimilarity (1-Spearman’s *R*) across all model response patterns for each layer output.

We then compared (Spearman’s *R*) the layer-specific deep scene model RDMs with the time-resolved MEG RDMs and averaged results over layers, yielding a time course indicating how well the deep scene model predicted and thus explained scene representations (Figure 3D). To compare against other models, we performed equivalent analyses to a deep neural network trained on object-categorization (termed deep object network) and standard models of object (HMAX) and scene-recognition (GIST) (Oliva and Torralba, 2001; Serre et al., 2007).

We found that the deep object and scene network performed similarly at predicting visual representations over time (Figure 3D, for details see Table 2A; for layer-resolved results see Supplementary Figure 5), and better than the HMAX and GIST models (for direct quantitative comparison see Supplementary Figure 6).

**Table 2.** Onset and peak latencies for model-MEG representational similarity analysis. Onset and peak latency *(n* = 15, *P* < 0.05, cluster-level corrected, cluster-definition threshold *P* < 0.05) with 95% confidence intervals. **A**) Correlation of models to MEG data. **B**) Comparison of MEG-model correlation for the deep scene network and all other models. 95% confidence intervals are reported in brackets.

| **A** |  |  |
| --- | --- | --- |
|  | **Onset latency** | **Peak latency** |
| GIST | 47 (45 - 149) | 80 (76 - 159) |
| HMAX | 48 (25 - 121) | 74 (61 - 80) |
| Deep object network | 55 (20 - 61) | 97 (83 – 117) |
| Deep scene network | 47 (23 - 59) | 83 (79 - 112) |
| **B** |  |  |
| Deep scene network minus GIST | 58 (50 - 78) | 108 (81 - 213) |
| Deep scene network minus HMAX | 75 (62 - 86) | 108 (97- 122) |
| Deep scene network minus deep object network | - | - |

**Table 3:**
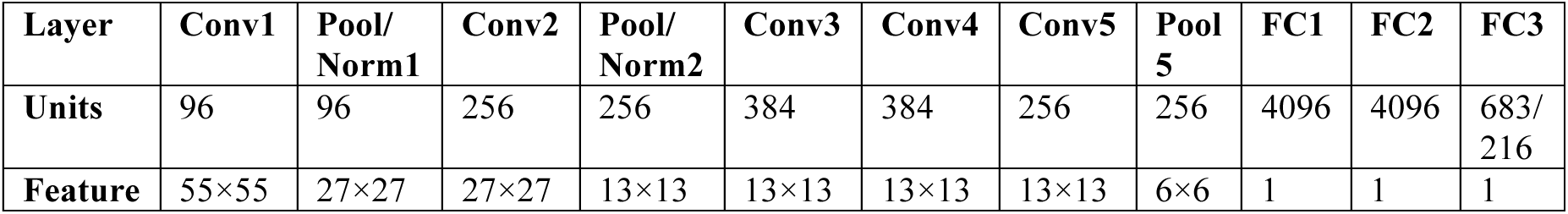
Number of units and features for each CNN layer. Units and features of the deep neural network architecture were similar as proposed in (Krizhevsky et al., 2012). All deep neural networks were identical with the exception of the number of nodes in the last layer (output layer) as dictated by the number of training categories, i.e. 683 for the deep object network, 216 for deep scene network. Abbreviations: Conv = Convolutional layer, Pool = Pooling layer; Norm = Normalization layer; FC1-3 = fully connected layers. The 8 layers referred to in the manuscript correspond to the convolution stage for layers 1-5, and the FC103 stage for layers 6-8 respectively.

In sum, our results show that brain representations of single scene images were best predicted by deep neural network models trained on real-world categorization tasks, demonstrating the ability of the models to capture the complexity of scene recognition, and their semblance to the human brain representations.

### 3.4 Representations of scene size emerged in the deep scene model

Beyond prediction of neural representations of single scene images, does the deep scene neural network indicate the spatial layout property scene size? To visualize, we used multidimensional scaling (MDS) on layer-specific model RDMs, and plotted the 48 scene images into the resulting 2D arrangement color-coded for scene size (black= small, gray = large). We found a progression in the representation of scene size in the deep scene network: low layers showed no structure, whereas high layers displayed a progressively clearer representation of scene size (A). A similar, but weaker progression, was visible for the deep object network (Figure 4B). Comparable analysis for HMAX and GIST (Figure 4C,D) found no prominent representation of size.

**Figure 4.**
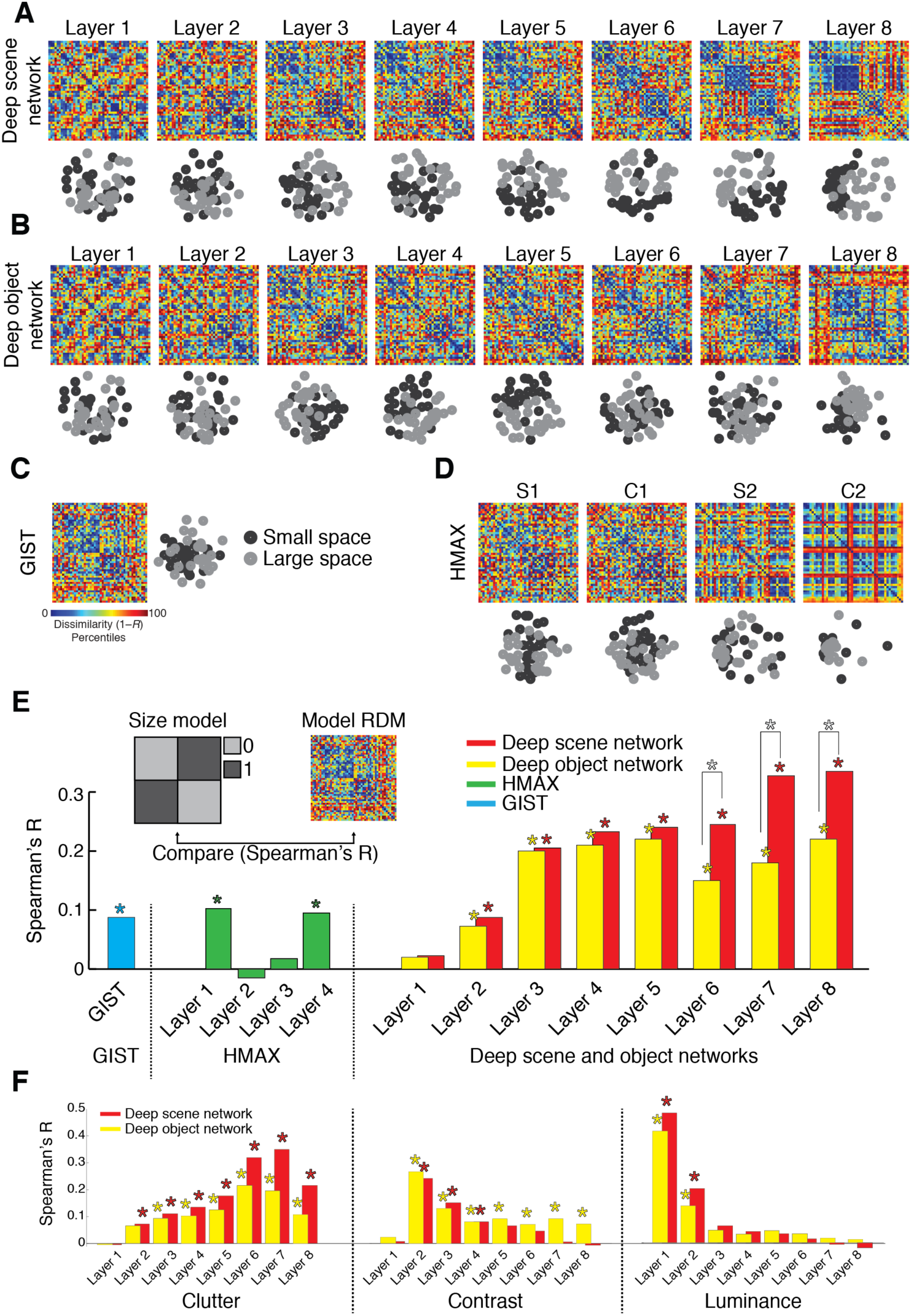
Representation of scene size in computational models of object and scene categorization. **A-D**) Layer-specific RDMs and corresponding 2D multidimensional scaling (MDS) plots for a deep scene network, deep object network, GIST, and HMAX. MDS plots are color-coded by scene size (small = black; large = gray). **E**) Quantifying the representation of scene size in computational models. We compared (Spearman’s *R)* each model’s RDMs with an explicit size model (RDM with entries 0 for images of similar size, 1 for images of dissimilar size). Results are color-coded for each model. **F**) Similar to (E) for clutter, contrast and luminance (results shown only for deep scene and object networks). While representations of the abstract scene properties size and clutter emerged with increasing layer number, the low-level image properties contrast and luminance successively abstracted away. Stars above bars indicate statistical significance. Stars between bars indicate significant differences between the corresponding layers of the deep scene vs. object network. Complete layer-wise comparisons available in Supplementary Figure 7. *(n* = 48; label permutation tests for statistical inference, *P* < 0.05, FDR-corrected for multiple comparisons).

We quantified this descriptive finding by computing the similarity of model RDMs with an explicit size model (an RDM with entries 0 for images of similar size, 1 for images of dissimilar size; Figure 4E inset). We found a significant effect of size in all models (*P* < 0.05, FDR-corrected, stars above bars indicate significance). The size effect was larger in the deep neural networks than in GIST and HMAX, it was more pronounced in the high layers, and the deep scene network displayed a significantly stronger effect of scene size than the deep object network in layers 6-8 (stars between bars; for all pair-wise layer-specific comparisons see Supplementary Figure 7). A supplementary partial correlation analysis confirmed that the effect of size in the deep scene network was not explained by correlation with the other experimental factors (Supplementary Figure 8).

Together, these results indicate the deep scene network captured scene size better than all other models, and that scene size representations emerge gradually in the deep neural network hierarchy. Thus representations of visual space can emerge intrinsically in neural networks constrained to perform visual scene categorization without being trained to do so directly.

### 3.5 Neural representations of scene size emerged in the deep scene model

The previous sections demonstrated that representations of scene size emerged in both neural signals (Figure 2) and computational models (Figure 4). To evaluate the overlap between these two representations, we combined representational similarity analysis with partial correlation analysis (Clarke and Tyler, 2014) (Figure 5A).

**Figure 5.**
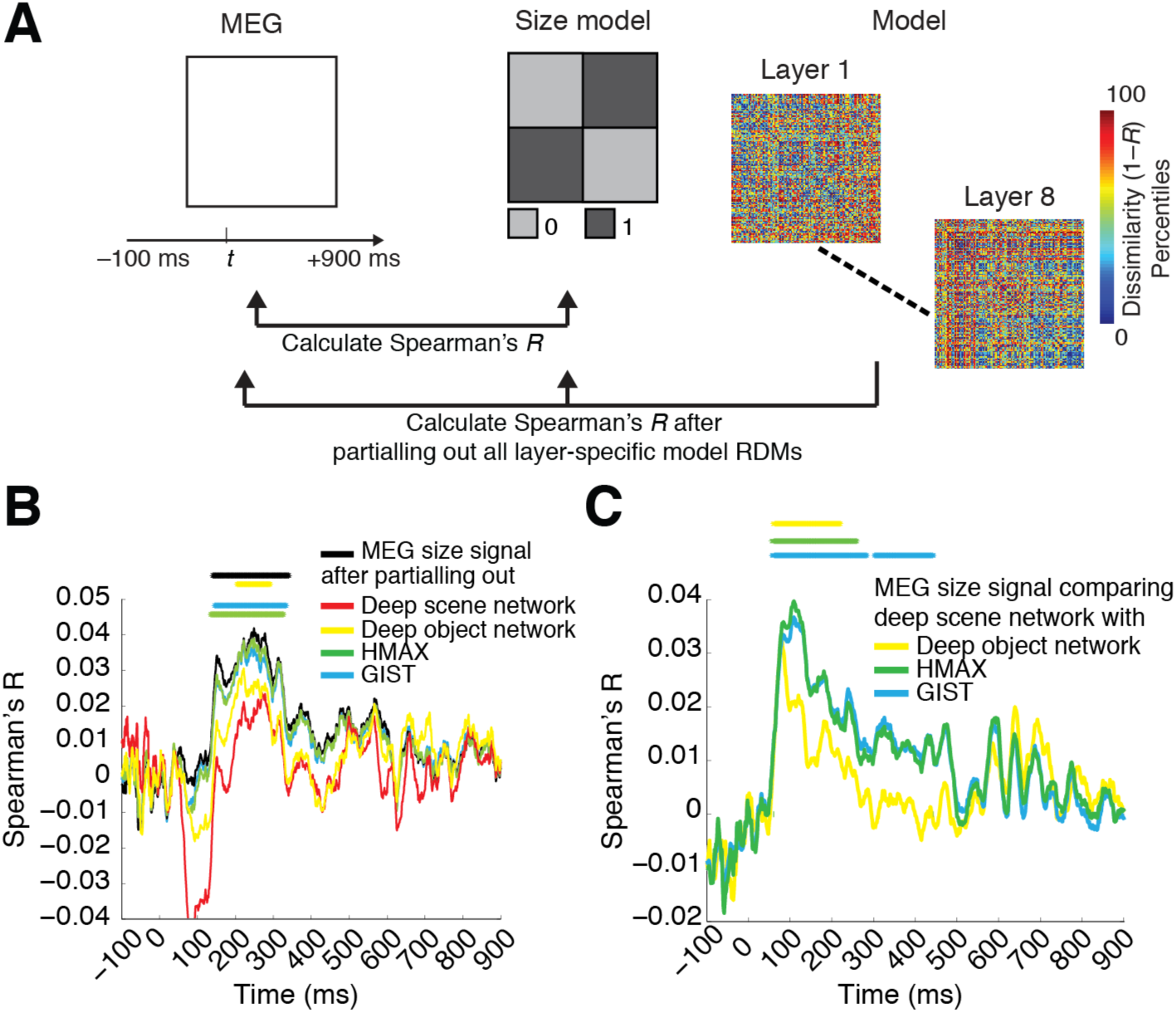
The deep scene model accounts for more of the MEG size signal than other models. **A**) We combined representational similarity with partial correlation analysis to determine which computational models explained emerging representations of scene size in the brain. **B**) MEG representations of scene size (termed MEG size signal) before (black) and after (color-coded by model) partialling out the effect of different computational models. Only partialling out the effect of the deep scene network abolished the MEG size signal. C) Difference in amount of variance partialled out from the size signal: comparing all models to the deep scene network. The deep scene network accounted for more MEG size signal than all other models *(n* = 15; cluster-definition threshold *P* < 0.05, significance threshold *P* < 0.05; results corrected for multiple comparisons by 5 for panel B and 3 for panel C).

We first computed the neural representations of scene size by correlating (Spearman’s *R*) the MEG RDMs with the explicit size model (black curve). We then repeated the process, but only after partialling out all layer-specific RDMs of a model from the explicit size model (color-coded by model) (Figure 5B). The reasoning is that if neural signals and computational models carry the same scene size information, the scene size effect will vanish in the latter case.

When partialling out the effect of the deep scene network, the scene size effect was considerably reduced and was no longer statistically significant. In all other models, the effect was reduced but was still statistically significant (Figure 5B). Further, the reduction of the size effect was higher for the deep scene network than all other models (Figure 5C). Equivalent analyses for scene clutter, contrast and luminance indicated that the deep scene and object networks abolished all effects, while other models did not (Supplementary Figure 9).

Together, these results show that only the deep scene model captured the neural representation of scene size in the human brain, singling it out as the best of the scene representation models tested here.

## 4 DISCUSSION

We characterized the emerging representation of scenes in the human brain using multivariate pattern classification methods (Carlson et al., 2013; Cichy et al., 2014) and representational similarity analysis (Kriegeskorte, 2008; Kriegeskorte and Kievit, 2013) on combined MEG and computational model data. We found that neural representations of individual scenes and the low-level image property contrast emerged early, followed by the scene layout property scene size at around 250 ms. The neural representation of scene size was robust to changes in viewing conditions and scene properties such as contrast, luminance, clutter level and category. Our results provide novel evidence for an electrophysiological signal of scene processing in humans that remained stable under real-world viewing conditions. To capture the complexity of scene processing in the brain by a computational model, we trained a deep convolutional neural network on scene classification. We found that the deep scene model predicted representations of scenes in the brain and accounted for abstract properties such as scene size and clutter level better than alternative computational models, while abstracting away low-level image properties such as luminance and contrast level.

### 4.1 A multivariate pattern classification signal for the processing of scene layout property scene size

A large body of evidence from neuropsychology, neuroimaging and invasive work in humans and monkeys has identified locally circumscribed cortical regions of the brain dedicated to the processing of three fundamental visual categories: faces, bodies and scenes (Allison et al., 1994; Kanwisher et al., 1997; Aguirre et al., 1998; Downing et al., 2001; Tsao et al., 2006; Kornblith et al., 2013). For faces and bodies, respective electrophysiological signals in humans have been identified (Allison et al., 1994; Bentin et al., 1996; Jeffreys, 1996; Liu et al., 2002; Stekelenburg and de Gelder, 2004; Thierry et al., 2006). However, electrophysiological markers for scene-specific processing have been identified for the auditory modality only (Fujiki et al., 2002; Tiitinen et al., 2006), and a visual scene-specific electrophysiological signal had not been described until now.

Our results provide the first evidence for an electrophysiological signal of visual scene size processing in humans. Multivariate pattern classification analysis on MEG data revealed early discrimination of single scene images (peak at 97ms) and the low-level image property contrast (peak at 74ms), whereas the abstract property of space size was discriminated later (peak at 249ms). While early scene-specific information in the MEG likely emerged from low-level visual areas such as V1 (Cichy et al., 2014), the subsequent scene size signal had properties commonly ascribed to higher stages of visual processing in ventral visual cortex: the representation of scene size was tolerant to changes occurring in real world viewing conditions, such as luminance, contrast, clutter level and category. The electrophysiological signal thus reflected scene size representations that could reliably be used for scene recognition in real world settings under changing viewing conditions (Poggio and Bizzi, 2004; DiCarlo and Cox, 2007; DiCarlo et al., 2012). This result paves the way to further studies of the representational format of scenes in the brain, e.g. by measuring the modulation of the scene-specific signal by other experimental factors.

The magnitude of the scene size effect, although consistent across subjects and statistically robust to multiple comparison correction, is small with a maximum of ~1%. Note however that the size effect, in contrast to single image decoding (peak decodability at ~79%), is not a measure of how well single images differing in size can be discriminated, but a difference measure of how much better images of different size can be discriminated rather than images of the same size. Thus, it is a measure of information about scene size over-and-above information distinguishing between any two single scenes. The magnitude of the size effect is comparable to effects reported for abstract visual properties such as animacy (1.9 and 1.1% respectively, Cichy et al., 2014).

What might be the exact locus of the observed scene size signal in the brain? Previous research has indicated parametric encoding of scene size in parahippocampal place area (PPA) and retrosplenial cortex (Park et al., 2014), corroborating numerous studies showing that spatial properties of scenes such as boundaries and layout are represented in these cortical regions (Epstein and Kanwisher, 1998; Epstein et al., 1999; Wolbers et al., 2011b). Both onset and peak latency of the observed scene size signal concurred with reported latencies for parahippocampal cortex (Mormann et al., 2008), suggesting that one or several nodes of the human spatial navigation network might be the source of the scene size signal.

Last, we found that not only scene size representations, but also scene clutter representations were tolerant to changes in viewing conditions, and emerged later than the low-level image contrast representations. These results complement previous findings in object perception research that representations of single objects emerge earlier in time than representations of more abstract properties such as category membership (Carlson et al., 2013; Cichy et al., 2014).

### 4.2 Neural representations of abstract scene properties such as scene size are explained by a deep neural network model trained on scene classification

Scene processing in the brain is a complex process necessitating a formal quantitative model that addresses this complexity. Here, our study of several models of scene and object recognition provided three novel results, each with fundamental theoretical implications.

First, deep neural networks offered the best characterization of neural scene representations compared to other models tested. The superiority of high performing deep neural networks over simpler models indicates that hierarchical architectures might be necessary to capture the structure of single scene representations in the human brain. While previous research has established that deep neural networks capture object representations in human and monkey inferior temporal cortex well, we demonstrated that a deep neural network explained millisecond-resolved dynamics underlying scene recognition from processing of low- to high-level properties, better than other models of object and scene-processing tested. Concerning high-level abstract scene properties in particular, our results shed lights into the black box of cortical scene processing, providing novel insight both from the perspective of modeling, and of experimental brain science. From a modeling perspective, the near monotonic relationship between the representation of size and clutter level in the deep neural network and the network layer number indicates that scene size is an abstract scene property emerging through complex multi-step processing. From the perspective of experimental brain science, our results provide an advance in understanding neural representations of the processing of abstract scene properties such as spatial layout. Neuronal responses in high-level visual cortex are often sparse and nonlinear, making a full explanation by simple mathematical models in low-dimensional spaces or basic image statistics unlikely (Groen et al., 2013; Rice et al., 2014; Watson et al., 2014; Rice et al., 2014). Instead, our result concurs with the finding that complex deep neural networks performing well on visual categorization tasks represent visual stimuli similar to the human brain (Cadieu et al., 2013; Yamins et al., 2014), and extends the claim to abstract properties of visual stimuli.

The second novel finding is that a deep neural network trained specifically on scene categorization had superior representation of scene size compared to a deep neural network trained on objects. Importantly, it also offered the best account of neural representations of scene size in the MEG, indicating that the underlying algorithmic computations matched the neuronal computations in the human brain. This indicates that the constraints imposed by the task the network is trained on, i.e. object or scene categorization, critically influenced the represented features. This makes plausible the notion that spatial representations emerge naturally and intrinsically in neural networks performing scene categorization, such as in the human brain. It further suggests that separate processing streams in the brain for different visual content, such as scenes, objects or faces, might be the result of differential task constraints imposed by classification of the respective visual input (DiCarlo et al., 2012; Yamins et al., 2014).

The third novel finding is that representations of abstract scene properties (size, clutter level) emerged with increasing layers in deep neural networks, while low-level image properties (contrast, luminance) were increasingly abstracted away, mirroring the temporal processing sequence in the human brain: representations of low-level image properties emerged first, followed by representations of scene size and clutter level. This suggests common mechanisms in both and further strengthen the idea that deep neural networks are a promising model of the processing hierarchies constituting the human visual system, reinforcing the view of the visual brain as performing increasingly complex feature extraction over time (Thorpe et al., 1996; Liu et al., 2002; Reddy and Kanwisher, 2006; Serre et al., 2007; Kourtzi and Connor, 2011; DiCarlo et al., 2012).

However, we did not observe a relationship between layer-specific representations in the deep neural networks and temporal dynamics in the human brain. Instead, the MEG signal predominantly reflected representations in low neural network layers (Supplementary Figure 5). One reason for this might be that our particular image set differed strongly in low-level features, thus strongly activating early visual areas that are best modeled by low neural network layers. Activity in low-level visual cortex was thus very strong, potentially masking weaker activity in high-level visual cortex that is invariant to changes in low level features. Another reason might be that while early visual regions are close to the MEG sensors, creating strong MEG signals, scene-processing cortical regions such as PPA are deeply harbored in the brain, creating weaker MEG signals. Future studies using image sets optimized to drive low-and high level visual cortex equally are necessary, to test whether layer-specific representations in deep neural networks can be mapped in both time and in space onto processing stages in the human brain.

### 4.3 Conclusions

Using a combination of multivariate pattern classification and computational models to study the dynamics in neuronal representation of scenes, we identified a neural marker of spatial layout processing in the human brain, and showed that a deep neural network model of scene categorization explains representations of spatial layout better than other models. Our results pave the way to future studies investigating the temporal dynamics of spatial layout processing, and highlight deep hierarchical architectures as the best models for understanding visual scene representations in the human brain.

## 5 ACKNOWLEDGEMENTS

This work was funded by National Eye Institute grant EY020484 (to A.O.), National Science Foundation grant BCS-1134780 (to D.P.), McGovern Institute Neurotechnology Program (to A.O. and D.P.), a Humboldt Scholarship (to R.M.C), and was conducted at the Athinoula A. Martinos Imaging Center at the McGovern Institute for Brain Research, Massachusetts Institute of Technology. We thank Santani Teng for helpful comments on the manuscript.

